# A global metagenomic atlas uncovers ubiquitous biosynthetic potential linked to adaptation in extreme environments

**DOI:** 10.64898/2026.04.17.719132

**Authors:** Rubing Du, Ruolin He, Qi Qian, Zhen Li, Qiyong Tang, Zhidong Zhang, Xinming Xu, Haoran Peng, Jun Liu, Marnix H. Medema, Qing Xu

## Abstract

Extreme environments impose severe physicochemical stresses that drive microorganisms to evolve specialized survival strategies. Microbial secondary metabolites determined by biosynthetic gene clusters (BGCs) are recognized as important mediators of microbial adaptation to environmental stress. However, their ecological roles, particularly habitat-dependent preferences across different environments, remain poorly understood. Although extreme environments provide opportunities to mine microbiomes for unique adaptations, such research is hampered by a lack of systematic overview of its genomic diversity, BGC diversity, and the relationships between them. Here, we constructed a standardized extremophilic genomic catalogue (SEGC) from 1,462 metagenomic samples spanning seven representative extreme habitats. The catalogue comprised 54,661 metagenome-assembled genomes representing 21,805 species, 66.1% of which were previously uncharacterized. With this catalogue, we identified 162,855 BGCs distributed across 81.5% of MAGs. Gene cluster family analysis showed the strong habitat dependence largely explained by species-level habitat specificity. Terpene biosynthetic pathways illustrated habitat-linked adaptive strategies, with hopan-22-ol biosynthesis enriched in acid mine, deep sea and hydrothermal plume environments, while retinal-based phototrophy predominated in cryosphere and saline–alkaline habitats. Metatranscriptomic analyses supported *in situ* activity of these pathways. In conclusion, we presented a global atlas of biosynthetic potential across extreme-environment microbiota and revealed habitat-dependent patterns of secondary metabolism linked to microbial survival.

## 1. Introduction

Extreme environments, characterized by harsh physicochemical conditions such as extreme temperatures, salinity, and pH, are widely distributed on Earth ^1, 2^. Although these environments represent the ecological and evolutionary limits of life, diverse and numerous microorganisms have been found in every extreme environment, ranging from natural ecosystems, such as hot springs, to harsh settings caused by anthropogenic activities such as acid mine drainage ^3–5^. Recent studies have uncovered a rich diversity of microorganisms from all three domains of life in extreme environments, encompassing both lineages commonly detected in conventional habitats and environment-specific taxa ^4, 6^. To survive and thrive there, these microorganisms have evolved to retain or acquire specialized metabolic capabilities to cope with extreme environmental stresses ^7, 8^. Revealing these specialized metabolic strategies could help us understand how life handles harsh conditions and harness such strategies for biotechnological purposes.

In almost all ecosystems, microbial genomes contain biosynthetic gene clusters (BGCs) to synthesize various secondary metabolites, also known as specialized metabolites ^9, 10^. The major classes of secondary metabolites include ribosomally synthesized and posttranslationally modified peptides (RiPPs), polyketide synthases (PKSs), nonribosomal peptides (NRPSs), polyketide-nonribosomal peptide combinations (PKS-NRPS hybrids), and terpenes ^11, 12^. It has been well explored that various secondary metabolites possess antimicrobial properties and determine the outcome of microbial interactions ^13, 14^. Typically, microorganisms can synthesize antimicrobial peptides and use them as chemical weapons to kill or inhibit competitors and gain a competitive advantage in nutrients and space ^15, 16^. In addition, secondary metabolites also enable microbes to adapt to their abiotic environments ^17^.

For example, a few studies have revealed that certain taxa from Antarctic, such as *Flavobacteriaceae*, *Sphingobacterium* and *Hymenobacter*, synthesize carotenoids that support light-driven energy acquisition ^18^. Some specialized metabolites such as aryl polyenes could provide protection against oxidative stress ^19^, and the compatible solute ectoine acts as a protective agent against diverse environmental stresses ^20^. However, it remains unclear whether secondary metabolites function as a universal survival strategy, and whether distinct secondary metabolite production strategies show habitat-dependent preferences.

Despite the recognized importance of secondary metabolites, the majority of microbes on the planet are unculturable, representing a major bottleneck to uncovering their biosynthetic potential and ecological roles^21, 22^. Recent advances in metagenomics and genome-resolved analysis enable the reconstruction of metagenome-assembled genomes (MAGs) from uncultured taxa, substantially expanding the available genomic repertoire across diverse ecosystems^23^. While recent efforts have been made to reconstruct MAGs from extreme environment microbiomes ^24–26^, these studies were restricted by narrow sample coverage and limited geographical representation. Moreover, methodological inconsistencies across studies hinder the development of a standardized genomic catalogue through simple data merging. Therefore, a comprehensive global dataset is essential to systematically deciphering the roles of secondary metabolites in microbial adaptation to environmental stresses.

In this study, we aim to uncover the contribution of secondary metabolites to microbial adaptation across global extreme environments. To resolve the technical limitations, we established a standardized extremophilic genomic catalogue (SEGC), based on public sequence data as well as 115 newly generated metagenomes, comprising taxonomic groups distinct from those commonly found in normal ecosystems. Using SEGC, we revealed the biosynthetic diversity of secondary metabolites and uncovered habitat-dependent distribution patterns driven by taxonomic composition. We further characterized habitat-dependent preferences of secondary metabolic pathway and uncovered functional links between environmental features and microbial survival. Finally, we demonstrated that the lineages encoding above secondary metabolic pathways are broadly distributed in global extreme environments and unexpectedly enriched in novel species. Together, this work highlights the ecological importance of secondary metabolism in extremophiles and provides an accessible genomic resource for future research in ecology as well as resource mining.

## 2. Results

### 2.1 Global extreme environments harbor diverse and unique microbiota

To build a genomic catalogue of microorganisms from extreme environments, we conducted a global-scale analysis of 1,462 metagenomic samples with 37.44 Tb raw data (Fig. 1a) including1,347 publicly available samples and 115 in-house samples (see Methods for details). These combined datasets encompassed seven representative types of extreme environments, including acid mine, cryosphere, deep sea, hot spring, hydrothermal plume, saline-alkaline environment, and subsurface environment. Based on a standardized metagenomic per-sample binning pipeline, we recovered 195,841 preliminary MAGs, among which 54,661 MAGs met the medium-quality requirements ^27^, with an average completeness of 84.40% and an average contamination of 3.07% (Fig. 1b). Among these, 5,480 MAGs (10.03%) were classified as high-quality genomes. Collectively, these 54,661 MAGs constituted the SEGC, representing the largest genomic catalogue to date of global extreme-environment metagenomes.

**Fig. 1.**
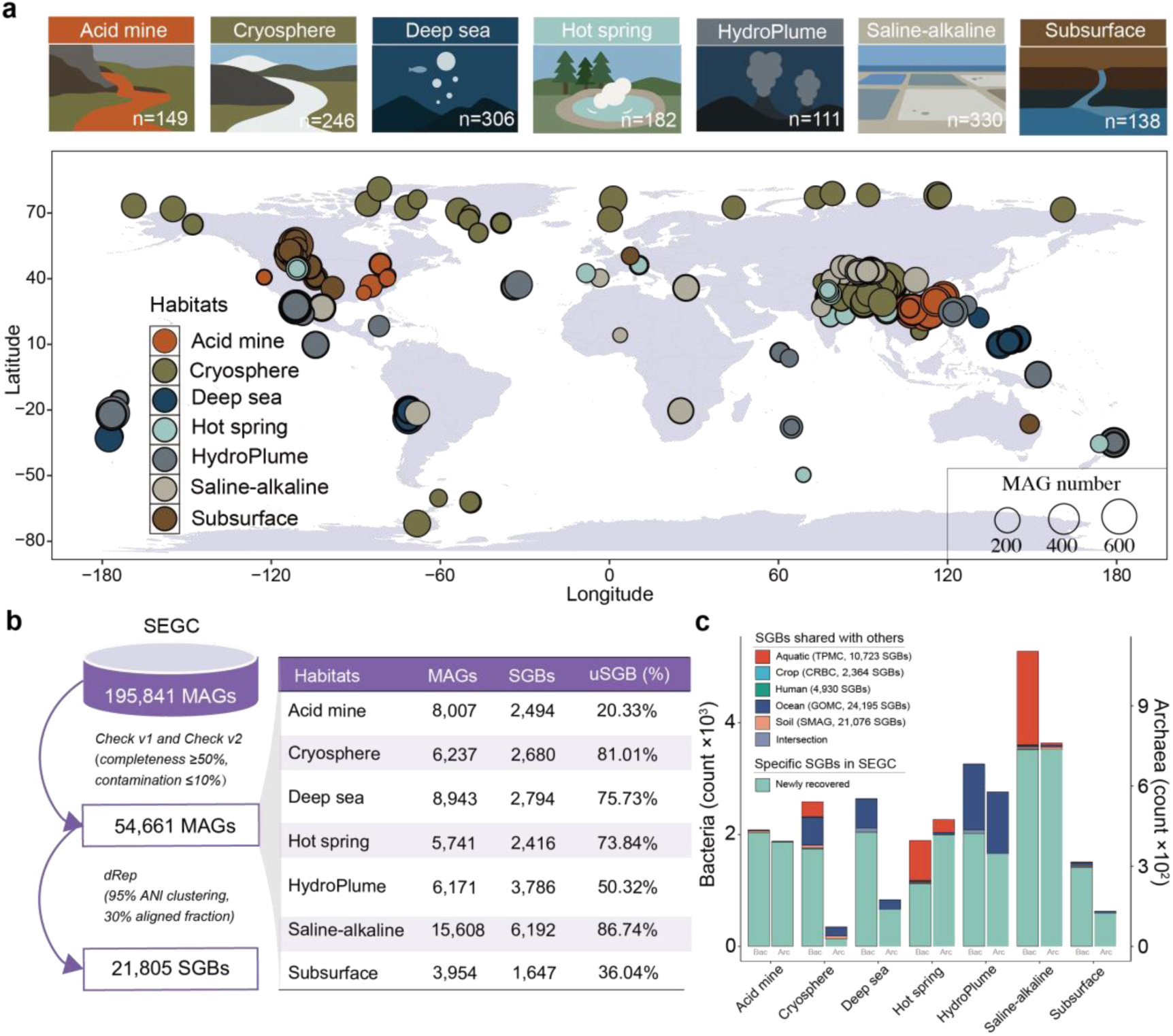
SEGC database. **(a)** The global distribution and statistics of samples and metagenome-assembled genomes (MAGs) recovered from each sample. Sample numbers from the seven extreme environments are shown within the corresponding habitat schematics. Dots on the world map indicate sampling locations, with dot size proportional to the number of MAGs recovered from each sample and colors denoting habitat types. HydroPlume represents hydrothermal plume. **(b)** Numbers of medium-quality MAGs recovered from each habitat. MAG quality was assessed using both CheckM1 and CheckM2, and medium-quality MAGs were de-replicated using dRep. The table summarizes, for each habitat, the numbers of MAGs and species-level genome bins (SGBs), as well as the proportion of unknown SGBs (uSGBs). **(c)** The uniqueness of SEGC compared to five common genomic datasets. Bar plots show the numbers of habitat-specific unique SGBs and shared SGBs. “Intersection” represents SGBs present in two or more non-SEGC datasets. “Bac” represents bacteria and “Arc” represents archaea.

We clustered all MAGs into 21,805 representative species-level genome bins (SGBs), including 19,229 bacterial species and 2,578 archaeal species (Fig. 1b). Comparisons with five representative public microbial genome databases, including aquatic ^28^, crop ^29^, human ^30^, ocean ^31^, and soil ^32^, demonstrated that our catalogue exhibits comparable species richness but a substantially higher archaeal representation (>10%), including 669 DPANN superphylum species. Cross-database comparisons showed that 16,022 species (73.48%) in our catalogue are unique to extreme environments (Fig. 1c).

Taxonomic annotation showed that 14,426 SGBs did not match any reference genomes (ANI < 95%) and were identified as unknown species (uSGBs). The uSGBs were detected in all sampled habitats, with their proportion ranging from 20.33% to 86.74%. The saline-alkaline and cryosphere environments harbored the highest proportions of uSGBs, each exceeding 80% (Fig. 1b). Among them, 3,664 SGBs could not be assigned to any known genus, 632 to a family, and 121 to an order according to the GTDB R220 taxonomy. These uSGBs spanned a wide taxonomic range, covering 92.00% of all detected phyla. Mapping metagenomic reads to SGBs revealed the relative abundance of uSGBs was significantly higher than that of known SGBs (kSGBs) (*P* =0.006), reaching more than twice that of kSGBs across 45.96% of samples. This disparity was most pronounced in cryosphere, saline-alkaline, and hot spring environments, where the proportion of reads mapping to uSGBs was up to ninefold higher than that for kSGBs. Among the 14,426 uSGBs, 1,546 were classified as archaea, expanding the number of archaeal species by 26.84%. Incorporating these uSGBs into phylogenetic analyses yielded a 17.74% increase in the total phylogenetic diversity across the life tree of archaea. Specifically, within the genera *PXRE01*, *Salinarchaeum*, *Halodesulfurarchaeum*, *SKNY01*, and *SKSH01*, the newly discovered species more than doubled the phylogenetic diversity. The genus *Salinarchaeum* previously contained three known species, and the SEGC contributed 15 additional species, resulting in a 149.30% increase in phylogenetic diversity.

### 2.2 Extreme-environment microbiota harbor broad and diverse biosynthetic potential

The SEGC provides a unique opportunity to explore biosynthetic potential across both known and previously uncharacterized species, spanning bacteria and archaea from global extreme environments. We predicted a total of 162,855 BGCs or BGC fragments from 44,546 MAGs (81.50%) distributed across all seven habitats, spanning 49.41% of archaeal SGBs and 86.04% of bacterial SGBs (Fig. 2a). These results suggest ubiquitous secondary metabolic potential in extreme-environment microbiota. These BGCs were categorized into eight classes (Fig. 2a), with terpenes (28.81%), RiPPs (25.73%) and NRPSs (12.61%) being the dominant classes. Among them, 72.23% of the identified BGCs were longer than 5 Kb, with a median length of 12.62 Kb. The largest BGC was found in a cryosphere-derived microorganism (phylum Bacillota_A, genus *JAEZXR01*), encoding a remarkable number of 22 PKS modules. Notably, 61.46% of bacterial BGCs and 50.86% of archaeal BGCs originated from uSGBs.

**Fig. 2.**
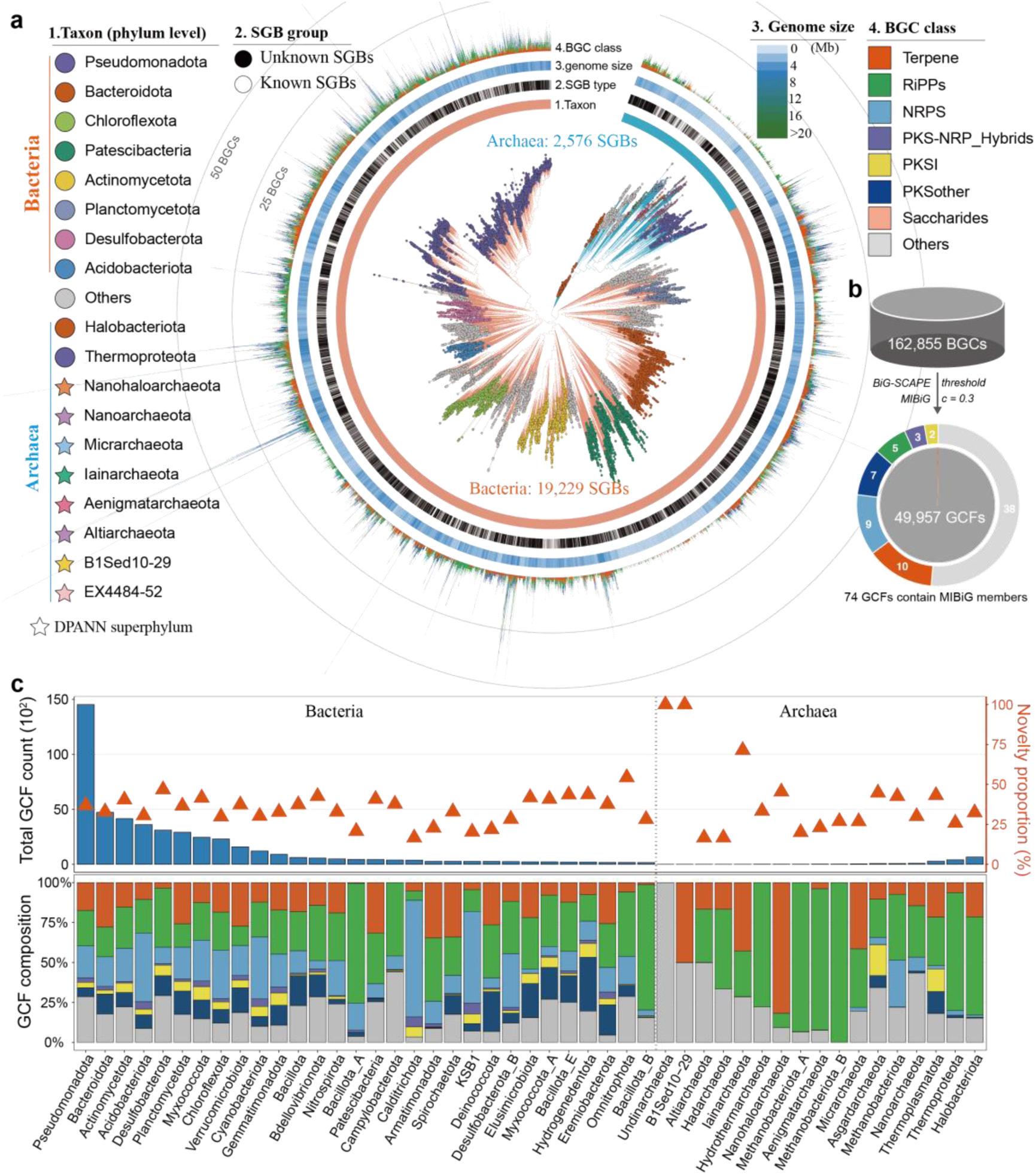
Biosynthetic potential of extreme-environment microbiota. **(a)** Biosynthetic potential of archaea and bacteria. Archaeal and bacterial phylogenetic trees were manually combined, with branch colors distinguishing archaea and bacteria and terminal nodes colored by phylum. Concentric annotations from inner to outer rings indicate taxonomic assignment, SGB type, genome size, and biosynthetic gene cluster (BGC) class. **(b)** Diversity of BGCs. A total of 162,855 BGCs were clustered into 49,957 gene cluster families (GCFs) using BiG-SCAPE. Among them, 74 GCFs correspond to characterized BGCs in the MIBiG database, with their classifications and numbers shown in the outer ring of the pie chart. **(c)** Novelty of microbial biosynthetic potential. Bar plots indicate the number of GCFs identified in each phylum, while triangular points show the proportion of novel GCFs compared to the BiG-FAM database. Stacked bars represent the distribution of GCF classes across phyla. For bacteria, the top 30 phyla with the highest GCF numbers are shown, whereas all archaeal phyla are included.

The BGC numbers varied widely across species, ranging from 1 to 175 BGCs per species. Among 162,855 BGCs, only 17,239 were annotated as complete, which is consistent with previous studies ^33, 34^. To reduce fragmentation bias, we focused on BGCs longer than 5 kb to analyze phylogenetic patterns of biosynthetic potential according to previous studies ^9, 35^. Bacteria possessed greater biosynthetic potential than archaea in extreme environments, supported by higher BGC counts (Wilcoxon *P* = 3.45e−127; |Cohen’s d| = 0.53). Within archaea, Halobacteriota exhibited the highest metabolic potential, whereas members of the DPANN superphylum encoded very few BGCs (Kruskal-Wallis chi-squared = 222.44, P = 3.13e−45). In bacteria, we observed clade-specific enrichment, with taxa such as Acidobacteriota, Nitrospirota_A and Desulfobacterota_B exhibiting markedly high biosynthetic potential, whereas Patescibacteria displayed very low levels (Kruskal-Wallis chi-squared = 1841.5, *P* < 2.2e-16). Meanwhile, compared with bacterial BGCs, archaeal BGCs were skewed toward RiPPs (49.47%) and terpenes (27.09%), whereas NRPS (1.43%) and PKS (4.75%) pathways were largely absent, suggesting evolutionary constraints on archaeal secondary metabolism.

To examine habitat-dependent preferences in biosynthetic potential, we compared BGC number and composition across different environments. Microorganisms from each habitat displayed comparable levels of biosynthetic capacity (2.54-3.71 BGCs per species). Microorganisms from the cryosphere showed the highest biosynthetic potential, whereas microorganism from the hot springs exhibited the lowest biosynthetic potential (Kruskal−Wallis *P* = 1.14e−92). In terms of BGC composition, we found cryosphere and saline-alkaline habitats to exhibit a higher biosynthetic potential of terpenes (Wilcoxon *P* <0.05). This pattern suggests that cryosphere and saline-alkaline microorganisms may rely more heavily on terpene secondary metabolism for ecological adaptation, although the biological roles of these pathways remain unclear.

BGC clustering analysis showed that 162,855 BGCs were grouped into 49,957 GCFs, including 12,741 RiPP, 8,494 terpene, and 10,780 NRPS GCFs. Comparison with MIBiG revealed that only 74 GCFs matched known products (Fig. 2b), leaving 99.85% of GCFs uncharacterized. These results highlight the enormous unexplored biosynthetic landscape. The complete BGCs belonged to 6,874 GCFs and 6,841 GCFs lacked MIBiG annotation, including 2,141 RiPP, 1464 terpene, and 519 NRPS GCFs. BiG-FAM catalogs 1,225,071 BGCs from 209,206 genomes and MAGs, and comparison with this reference showed that 18,650 GCFs in our dataset display no or low similarity to BiG-FAM BGCs (BiG-SLiCE distances >900), the latter being from isolate genomes and non-extreme environments. Altogether, these results suggest that BGCs in our dataset are likely to encode production of novel molecules or production of known molecules for which no biosynthetic pathway has yet been characterized.

These novel GCFs were found not only in newly discovered species but also in known species from typical prolific phyla such as Pseudomonadota, Bacteroidota, and Actinomycetota (Fig. 2c).

### 2.3 Taxonomic differences drive habitat-dependent biosynthetic potential

Although all eight GCF types were detected across extreme habitats (Fig. 3a), only 4,156 (8.32%) of GCFs were shared in more than one habitat (Fig. 3b and c). This pattern persisted even under a more relaxed similarity threshold (GCC level, c = 0.7), with 96.01% of GCCs remaining habitat-dependent, indicating substantial divergence in secondary metabolic pathways between environments. The largest number of shared GCFs (505 GCFs) occurred between cryosphere and saline–alkaline habitats, including 205 terpene, 89 RiPP, and 75 PKSother GCFs. Meanwhile, only 25 GCFs were shared across all habitats, and all of these GCFs belonged to RiPP pathways.

**Fig. 3.**
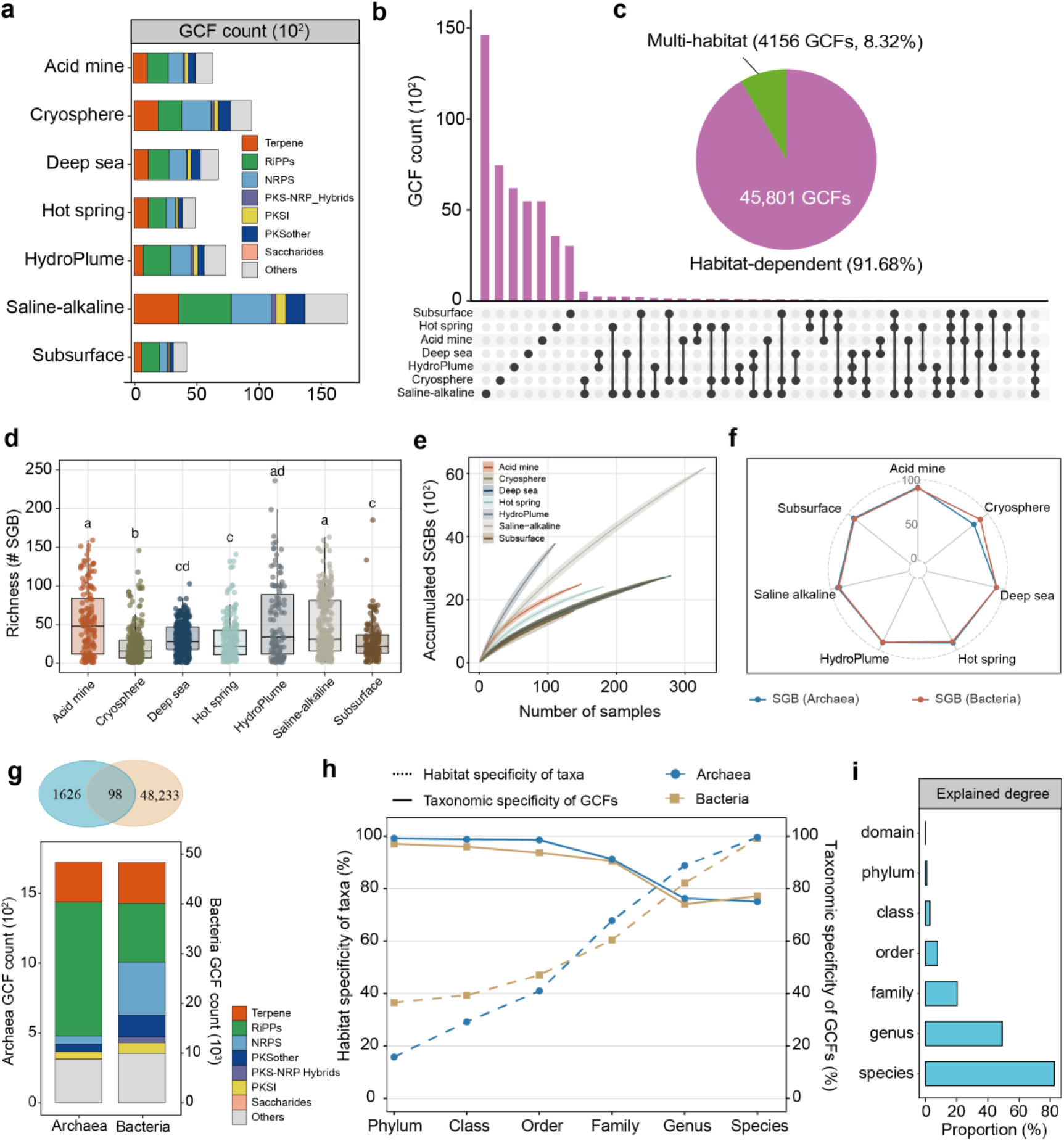
Habitat-dependent biosynthetic potential of extreme-environment microbiota. **(a)** Diversity of biosynthetic potential across different habitats. Stacked bar charts show the composition and abundance of gene cluster families (GCFs) in each habitat. HydroPlume represents hydrothermal plume. **(b)** Distribution patterns of GCFs across habitats. An UpSet analysis illustrates the distribution of habitat-dependent and shared GCFs among different habitats. **(c)** Numbers and ratios of habitat-dependent and multi-habitat GCFs. **(d)** Species-level diversity of SGBs across habitats. Each dot represents an individual sample. Letters indicate statistically significant differences of species richness among habitats (*P* < 0.05). **(e)** Species accumulation curves as a function of sample number for different habitats. **(f)** Habitat dependence of species at the SGB level, with archaea and bacteria distinguished by different colors. **(g)** Domain-level specificity of biosynthetic potential in bacteria and archaea. The Venn diagram shows the number of GCFs shared between bacteria and archaea, while stacked bar charts display the number and class composition of GCFs in each domain. **(h)** Ratio of species with habitat specificity (dashed lines) and GCFs with taxonomic specificity (solid lines) at different taxonomic levels. Archaea are shown in blue, whereas bacteria are shown in yellow. **(i)** Contribution of habitat-specific taxa to habitat-dependent GCFs. The explained proportion indicates the fraction of habitat-dependent GCFs that are derived from habitat-dependent species and GCFs are specific to those species.

Habitat-associated variation in GCFs may arise from taxonomic turnover between environments. We next assessed SGB distributions across habitats. SGB richness varied widely, with acid mine, hydrothermal plume, and saline-alkaline ecosystems showing the greatest diversity (Fig. 3d). By contrast, cryosphere microbiota showed unexpectedly lower richness (*P* <0.05) despite the highest biosynthetic capacity in this habitat. Species accumulation curves did not reach saturation even with increased sampling, indicating that microbial diversity in extreme environments remains substantially underestimated (Fig. 3e). Species distributions were strongly habitat-dependent, and this pattern was consistent across both bacteria and archaea (Fig. 3f). At broader taxonomic scales, more than 80% of genera and over 60% of families were restricted to single habitats (Fig. 3h).

To assess whether this taxonomic difference contributes to habitat-dependent biosynthetic profiles, we investigated the phylogenetic distribution of GCFs. Cross-domain comparisons revealed that only 98 GCFs were shared between bacteria and archaea (Fig. 3g). At finer taxonomic levels, GCFs remained highly lineage-specific; although specificity decreased at the genus and species levels, more than 70% of GCFs exhibited clear taxonomic limitations (Fig. 3f). We then quantified the extent to which habitat-specific GCFs could be explained by originating from habitat-specific taxa. More than 50% of habitat-specific GCFs originated from taxa restricted to the corresponding habitat at the genus level, and this relationship was strongest at the species level, where 81.50% of habitat-specific GCFs originated from species with habitat-restricted distributions (Fig. 3i). These results indicate that species-level taxonomic dependence of habitat gives rise to the pronounced habitat dependence observed in biosynthetic potential.

### 2.4 Biosynthetic potential displays habitat preferences linked to survival strategies

To reveal habitat-dependent biosynthetic potential, we analyzed structural domain preferences across environments using phylum- and habitat-stratified GCF dereplication. After filtering, 39,781 non-redundant BGC representatives (>5 kb) were retained. Comparative analysis of domain occurrence within each phylum revealed widespread habitat-specific enrichment patterns, with terpene-associated domains dominating the most significant signals (Fig. 4a). In detail, among a total of 406 domain categories showing highly significant enrichment (*P* < 0.001), 153 were associated with terpene BGCs, followed by 72 and 45 domains derived from RiPP and NRPS BGCs, respectively (Fig. 4a). Notably, 20 domains showed extremely strong enrichment (*P* < 10⁻²⁰), all derived from terpene BGCs, indicating pronounced habitat specificity of terpene biosynthesis. These domains were primarily affiliated with Actinomycetota and Pseudomonadota. In Actinomycetota, key enriched domains included squalene-hopene cyclase (PF13243, PF13249) and radical SAM enzymes (PF04055), which are involved in triterpenoid scaffold formation and were most prevalent in acid mine environments. In contrast, Pseudomonadota showed strong enrichment of domains associated with retinal-based phototrophy, including lycopene cyclase (PF05834), β-carotene dioxygenase (PF15461), and bacteriorhodopsin-like proteins (PF01036), particularly in cryosphere and saline–alkaline habitats.

**Fig. 4.**
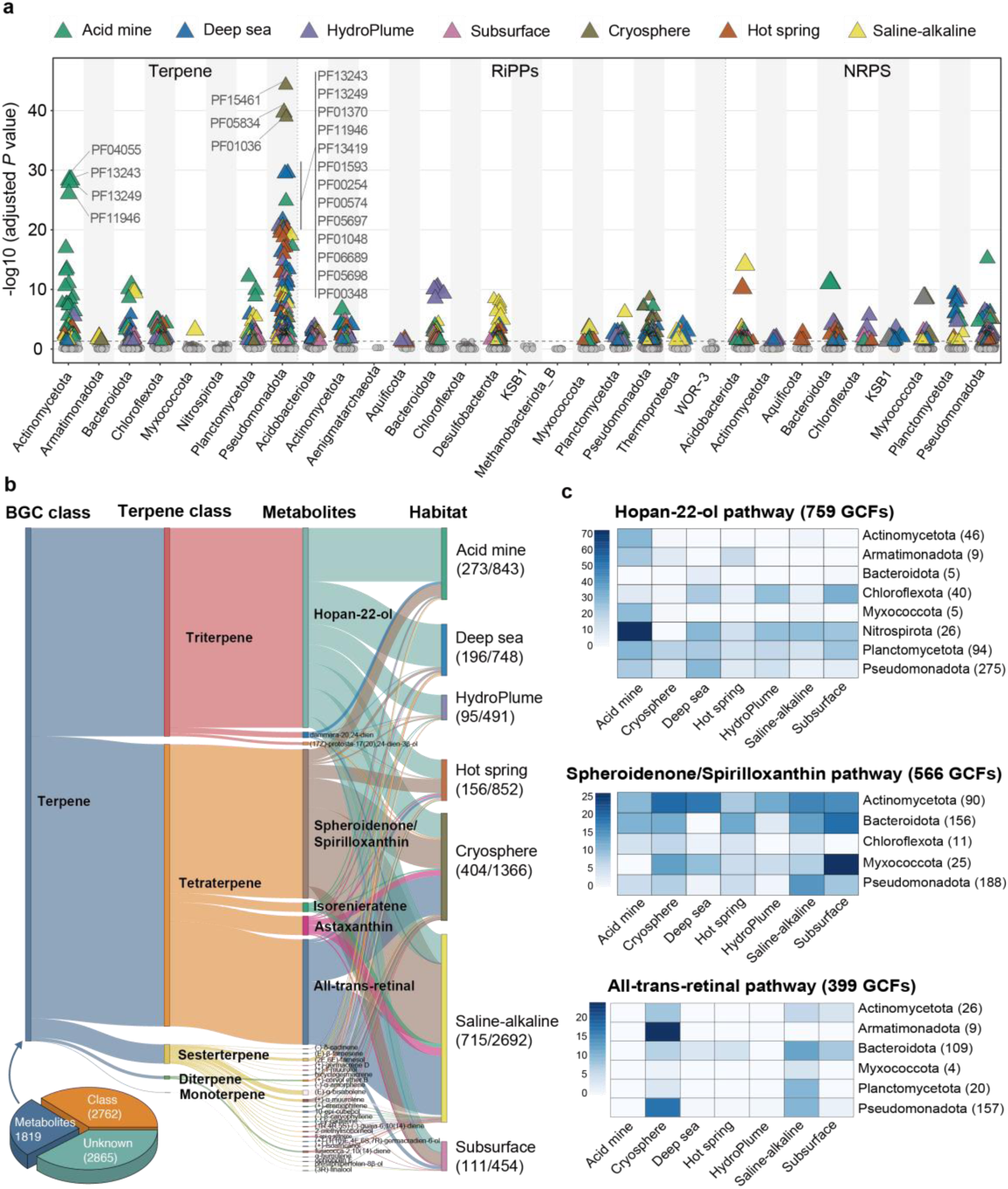
Habitat preferences in biosynthetic potential of secondary metabolites. **(a)** Habitat preferences of structural domains. A Manhattan plot shows habitat-associated enrichment of structural domains. BGCs were de-replicated based on gene cluster family (GCF), habitat, and phylum to minimize redundancy. *P* values indicate differences in domain occurrence frequencies across habitats. Each triangle represents a domain and is colored according to the habitat in which it appears most frequently. HydroPlume represents hydrothermal plume. **(b)** Diversity of terpene biosynthetic pathways. The three-dimensional pie chart (lower left) shows the numbers and ratio of terpene GCFs that could be assigned at the class and metabolite level. Terpene classes and metabolite categories were determined based on pathway-specific marker genes. Clusters were assigned as “Unknown” when no marker gene was detected in the BGC. The 1,819 terpene BGCs that can be assigned at the metabolite level are further associated with habitat. Numbers shown below each habitat indicate the counts of terpene GCFs that could be classified at the class level and the total number of terpene GCFs identified in that habitat. **(c)** Distribution frequency of three terpene biosynthetic pathways across taxonomic groups. The heatmap shows the distribution frequency of each pathway across different phyla and habitats, with numbers in parentheses indicating the numbers of GCFs corresponding to each metabolite pathway.

To further resolve functional preferences, we reconstructed terpene biosynthetic pathways based on EC annotation of non-redundant BGCs. A total of 4,581 BGCs was classified into defined biosynthetic pathways based on marker genes corresponding to the biosynthesis of distinct terpene skeletons. Of the classified clusters, 55.42% were assigned to triterpene pathways and 36.77% to tetraterpene pathways. Triterpenes were mainly derived from the hopan-22-ol pathway, which is associated with adaptation to low-oxygen ^36^ and low-pH conditions ^37^, and showed the highest occurrence in acid mine, followed by deep sea and hydrothermal plume environments. In contrast, tetraterpenes were dominated by spheroidenone/spirilloxanthin and retinal pathways, which are linked to photoprotection and light-driven energy acquisition. These pathways were preferentially enriched in cryosphere and saline–alkaline habitats and were largely absent from deep sea environments.

Consistent patterns between domain-level enrichment and pathway reconstruction highlight that terpene biosynthetic strategies are tightly coupled to environmental selection pressures. Specifically, hopanoid biosynthesis supports adaptation to oxygen- and pH-limited conditions, whereas carotenoid and retinal pathways are favored in light-exposed habitats (Fig. 5), reflecting distinct ecological strategies across extreme environments.

**Fig. 5.**
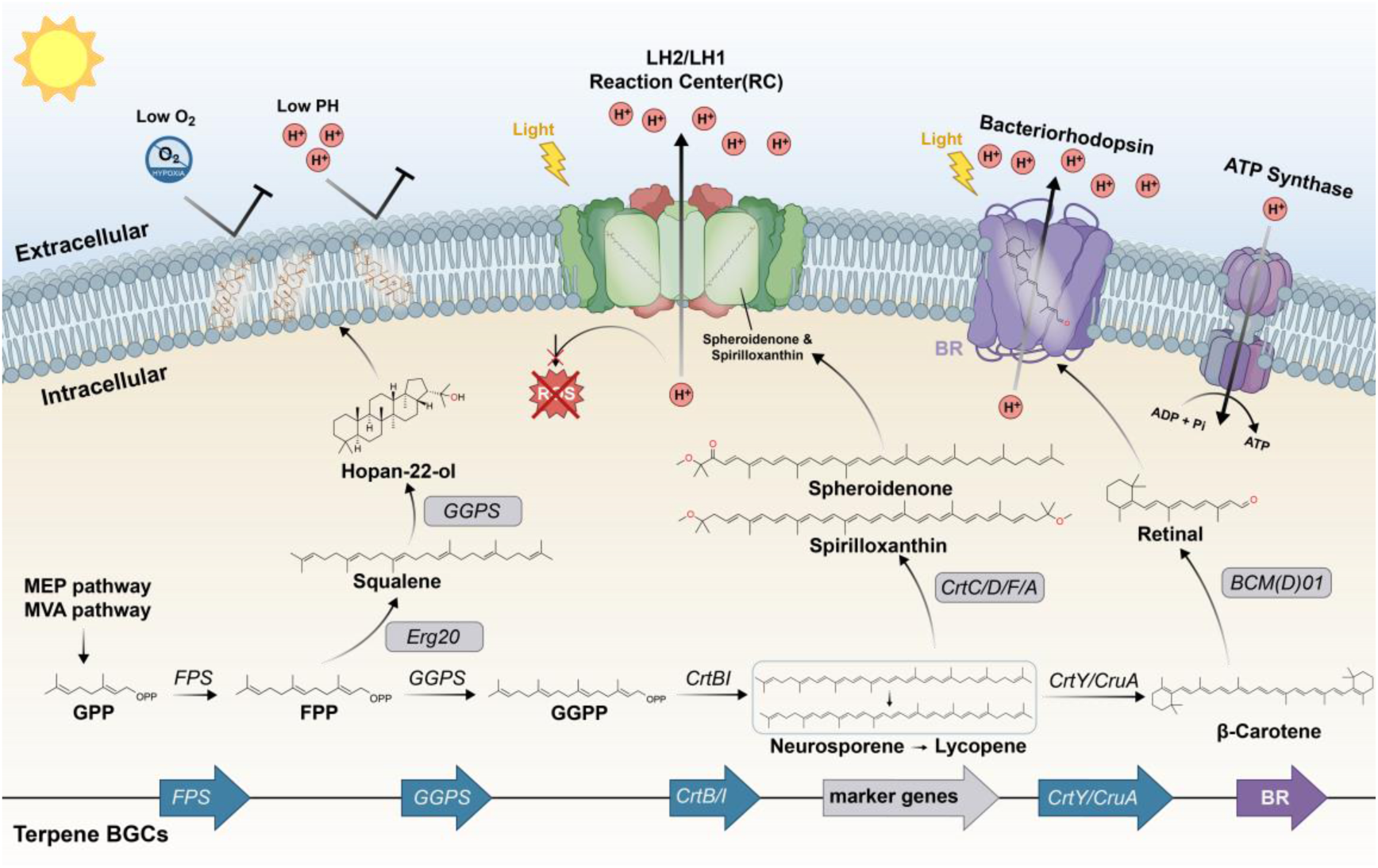
Terpene-mediated survival mechanisms in extremophiles. The bottom panel illustrates the biosynthetic pathways of three representative terpene-derived metabolites. Genes shown in blue represent common biosynthetic genes in terpene BGCs, grey indicates pathway-specific marker genes, and purple denotes bacteriorhodopsin-like proteins. Schematic diagrams illustrate survival strategies mediated by hopan-22-ol, spheroidenone/spirilloxanthin, and retinal. Hopan-22-ol is embedded in the cell membrane, where it enhances membrane stability and helps microorganisms tolerate low-oxygen and low-pH stress. Spheroidenone and spirilloxanthin participate in light-harvesting processes by capturing wavelengths not absorbed by chlorophyll and simultaneously protect cells against reactive oxygen species (ROS). Retinal, together with bacteriorhodopsin-like proteins, forms a retinal-based phototrophic system in the cell membrane. During light-driven proton pumping, protons are expelled from the cytoplasm, generating a proton motive force that subsequently drives ATP synthesis via ATP synthase.

### 2.5 Extreme environments retain ancient light-driven energy systems with functional redundancy

We then investigated the geographical distribution of retinal-based photosynthetic system in extreme ecosystems. All terpene BGCs (46,913) were screened for the presence of β-carotene dioxygenase (EC:1.13.11.63). A single-copy β-carotene dioxygenase was identified in 3,497 BGCs, originating from 332 samples. Among them, 299 samples were derived from cryosphere and saline–alkaline environments, with an average of 10 gene copies per sample (Fig. 6a). These samples collectively represented nearly 50% of all cryosphere and saline–alkaline samples analyzed (Fig. 6b). This indicates a broadly distributed, light-driven survival strategy in these extreme ecosystems. In cryosphere and saline–alkaline habitats, this function was often maintained by complex microbial communities rather than a single species. For example, more than 10 microbial species contributed to the retinal pathway in over 144 samples, and 72 samples harbored more than 20 contributing species. Notably, novel species accounted for an average of 77.84% of the β-carotene dioxygenase genes, highlighting their dominant contribution to this light-driven process and underscoring the ecological importance of newly discovered taxa in extreme environments (Fig. 6c).

**Fig. 6.**
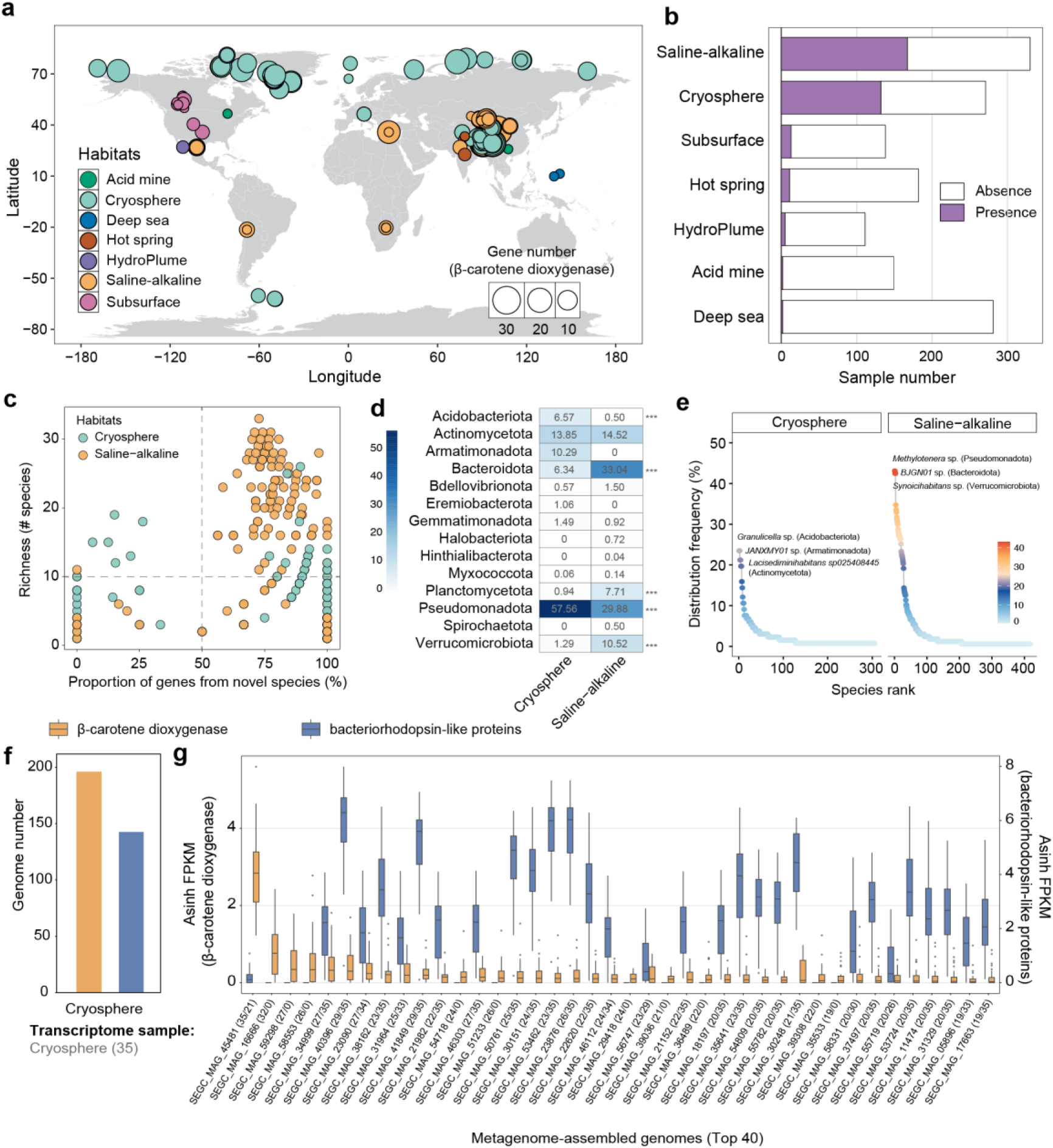
Global distribution of retinal biosynthesis pathways across extreme environments. **(a)** Geographic distribution of samples encoding the retinal biosynthesis pathway. Dots on the world map indicate sampling locations, with dot size proportional to the number of β-carotene dioxygenase genes detected from each sample and colors denoting habitat types. HydroPlume represents hydrothermal plume. **(b)** Proportion of samples containing the retinal biosynthesis pathway across different habitats. **(c)** Species diversity associated with the retinal biosynthesis pathway within samples and the relative contribution of novel taxa. **(d)** Major microbial contributors at the phylum level, with values indicating their relative contribution to retinal biosynthesis potential. **(e)** Distribution frequency of species harboring retinal biosynthesis pathways among samples in the corresponding habitats. **(f)** Numbers of transcriptionally active genomes encoding β-carotene dioxygenase and bacteriorhodopsin-like proteins detected in cryosphere environments. **(g)** MAGs exhibiting high transcriptional activity of retinal biosynthesis genes and bacteriorhodopsin-like proteins in polar environments (top 40). Numbers in parentheses following each MAG ID indicate the number of samples in which transcripts of retinal biosynthesis genes and bacteriorhodopsin-like proteins were detected.

The retinal-based phototrophic species exhibited remarkable phylogenetic diversity in both cryosphere and saline–alkaline habitats. These species spanned 14 phyla, 30 classes, 75 orders, 134 families, and 303 genera. Based on gene number, these taxa were primarily enriched in Pseudomonadota, Bacteroidota, Verrucomicrobiota, Planctomycetota, and Actinomycetota phyla. In saline–alkaline habitat, the genes were predominantly found in genomes of members of the Pseudomonadota phylum, whereas in the cryosphere they were found across both Pseudomonadota and Bacteroidota phyla (Fig. 6d). Interestingly, several of these phototrophic lineages have not been previously reported, revealing previously unrecognized diversity of light-utilizing microorganisms in extreme environments. We further investigated whether any species were prevalent carriers of this retinal-based system. We calculated the distribution frequency of functional species within the same habitat, and observed pronounced inter-sample variability in the taxonomic composition at species level (Fig. 6e). In cryosphere, the distribution frequency of all species was below 30%, with only four species exceeding 20% frequency. In saline–alkaline habitat, the median frequency was 0.75%, and only four species exhibited frequencies higher than 40%.

These results indicate that extreme environments maintain high sequence-level diversity but strong functional redundancy, with similar ecological functions fulfilled by distinct microbial lineages across samples.

To demonstrate whether these genes are transcribed *in situ*, we analyzed metatranscriptomic datasets to assess their transcriptional activity. Among 246 cryosphere and 330 saline–alkaline samples, 35 cryosphere samples had corresponding metatranscriptomes. These 35 samples, part of the Tara Oceans expedition, originated from the Arctic Ocean. Within these samples, 196 of 943 cryosphere MAGs exhibited clearly detectable transcriptional activity (presented in ≥2 samples) of β-carotene dioxygenase genes. Most MAGs exhibited a negative correlation between transcriptional activity and sequencing depth, with 22 MAGs showing statistically significant associations (Spearman FDR *P*< 0.05). Since retinal functions together with bacteriorhodopsin-like proteins (Fig. 5), we further investigated the genomic co-occurrence and transcriptional activity of bacteriorhodopsin-like proteins within BGCs. Overall, 69.10% of BGCs harboring β-carotene dioxygenase also encoded bacteriorhodopsin-like proteins. Among the 196 MAGs from cryosphere environments in which β-carotene dioxygenase was transcriptionally active, 149 MAGs exhibited concurrent expression of bacteriorhodopsin-like proteins. Consistent co-transcription of these two genes was detected across all 35 Arctic Ocean samples. Taxonomically, these MAGs spanned four phyla, dominated by Pseudomonadota (177 MAGs), followed by Actinomycetota (10 MAGs), Bacteroidota (6 MAGs), and Verrucomicrobiota (3 MAGs). These results expand the known diversity of phototrophic microorganisms in the Arctic Ocean ^38^ and reveal the co-occurrence and co-expression of retinal biosynthesis genes and bacteriorhodopsin within the same gene clusters, indicating the evolutionary conservation of this light-driven functional unit.

## 3. Discussion

The limited cultivability of microorganisms from extreme environments has long hindered comprehensive exploration of microbial life in extreme environments. Here, we present the largest comprehensive genomic catalogue of extreme environment microbiomes (SEGC), constructed using metagenomic binning approaches. Compared to previously representative studies, such as the genome recovery of 475 SGBs from 152 hot spring samples ^25^, 526 SGBs from 41 Arctic seawater samples ^24^, and 968 SGBs from glacier microbiomes ^26^, our study significantly expands the publicly available database of microbial genomes from extreme environments, covering seven representative extreme environments with broad geographical distribution. Using this extensive catalogue, we highlight that such environments are important archaeal repositories. The SEGC expands the life tree of archaea by 17.74% and includes a wide variety of DPANN archaea, which are considered representatives of the microbial “dark matter” ^39^. Meanwhile, we identified 21,805 prokaryotic species and uncovered strongly habitat-dependent distribution patterns. These results also imply a substantial underestimation of microbial diversity in global extreme environments, as further evidenced by the rarefaction curves based on current sequencing data, which could be addressed by increasing both sampling density and environmental coverage in future studies.

The SEGC catalogue demonstrates the diverse biosynthetic potential of extreme environment microbiota. Notably, the lion’s share of these BGCs and GCFs are significantly different to the known products in the MIBiG database and showed a high degree of diversity at the species level, which underscores extremophiles as a vast and untapped reservoir of secondary metabolites. In the BGC class composition, SEGC resembled ocean microbiomes, both dominated by terpene biosynthetic pathways ^31^. Distinct patterns have also been observed in other ecosystems. For example, the soil microbiome was enriched in NRPSs (23.8%) ^32^, the human-associated microbiome was dominated by RiPPs (74.5%) ^40^. These differences in BGC composition could guide the targeted genome mining to find the desired classes of secondary metabolites. However, most BGCs (89.41%) identified from the SEGC catalogue were fragmented, which hinders the discovery and characterization of novel secondary metabolites. It reflects limitations of short-read metagenomic sequencing, which is consistent with previous studies ^33, 34^. Emerging long-read sequencing platforms may be a solution to improve contig length and enable the recovery of a greater number of complete BGCs ^41, 42^. Even so, we have already identified 6,874 complete non-redundant BGCs in SEGC, including 2,141 RiPP, 1464 terpene, and 519 NRPS families, which are potential sources of novel antibiotics and bioactive compounds ^11^.

We also performed initial analyses of the ecological roles of secondary metabolism in extreme environments, such as microbial survival. Terpenes have long been considered chemical mediators of environmental adaptation ^43^. For example, previous studies have shown that rhodopsin-mediated phototrophy is widespread among marine planktonic bacteria ^44^ and has also been detected in archaea ^45, 46^. However, whether terpene-associated pathways constitute a universal microbial survival strategy, and whether their distribution reflects habitat-dependent preferences in global extreme environments, remains largely unresolved. Using this genomic catalogue, we highlight that terpenes exhibit both broad phylogenetic distribution and habitat-specific ecological preferences in extremophiles. In this study, we identified three widespread terpene biosynthetic pathways, including hopan-22-ol, spheroidenone/spirilloxanthin, and retinal biosynthetic pathways, that support microbial survival in extreme environments. These pathways exhibit clear habitat preferences that reflect distinct ecological pressures (Fig. 5). For example, in cryosphere and saline-alkaline habitats, where intense light reflection and strong physicochemical constraints limit carbon turnover, microbes enriched or retained retinal as well as chlorophyll (spheroidenone/spirilloxanthin)-based phototrophy. In contrast, acid mine, deep sea, and hydrothermal habitats favored the hopan-22-ol pathway, which is associated with membrane stabilization under low-oxygen ^36^ and low-pH conditions ^37^. By resolving the global distribution landscape of these pathways, this work provides new insights into how secondary metabolites mediate the interactions between microorganisms and extreme environments.

Complementary metatranscriptomic evidence supports the physiological importance of the retinal biosynthetic pathway in cryosphere. In particular, we detected transcription of the retinal biosynthetic pathway in all 35 Arctic Ocean samples. Although recently studies have revealed microbial adaptation by diverse carbon fixation strategies ^24, 47^, and metabolic reprogramming ^47, 48^, our results expand this knowledge, demonstrating that retinal-based phototrophy is a previously overlooked ecological trait contributing to microbial survival in the cryosphere. These findings also suggest that extreme environments may serve as a refuge for ancestral phototrophic mechanisms and provide a conceptual framework for re-evaluating the energy strategies of cryosphere microbiomes.

In summary, this study provides a large-scale and standardized genomic resource that maps the global biosynthetic landscape of extremophiles, substantially expanding known microbial diversity and biosynthetic potential. We demonstrate that secondary metabolites enhance microbial adaptability by facilitating energy acquisition and stress tolerance, and that their biosynthetic potential exhibits distinct habitat-encoded preferences. Together, these findings bridge biosynthetic potential and ecological function, and highlight the importance of secondary metabolism in the survival strategies of extremophiles.

## 4. Materials and methods

### Collection of in-house samples

The SEGC was developed based on 1,347 publicly available samples and 115 in-house samples. These 115 in-house soil samples were collected by the authors from representative saline–alkaline environments to improve the representation of this habitat type in the public datasets. These 115 soil samples were collected from four representative salt lakes (Daban Salt Lake, Ayding Lake, Huancai Lake, and Barkol Lake) and one abandoned salt mine (Qijiaojing) located in the northeastern Tianshan Mountains. Sampling points were selected based on visible soil salinization and positioned as close as possible to the shoreline, with sites spaced at least 1 km apart. In salt mining areas, samples were collected from salinized soils at 1-3 km intervals. The longitude, latitude, and elevation were measured using a portable GPS locator (Garmin, eTrex 201x, China). Soil was collected from a depth of 1-10 cm. For each sample, approximately 200 g of soil was retained after removing stones, plant debris, and salt blocks by sieving. All samples were immediately stored on dry ice and subsequently transferred to a -80 °C freezer until DNA extraction.

Water-soluble salts in the samples were quantified using the EC method. For each soil sample, 10 g of air-dried soil was extracted with 50 mL of deionized water (soil-to-water ratio 1:5, w/v) and shaken at room temperature for 30 min. The suspension was then filtered through quantitative filter paper. The EC of the filtrate was measured with a calibrated conductivity meter (DDSJ-308A, INESA Scientific Instrument, China). Soil pH was measured by mixing 10 g of dry soil with 10 mL of deionized water, shaking the mixture for 1 h, and then determining the pH with a pH meter (PHS-3C, INESA Scientific Instrument, China).

### DNA extraction and metagenomic sequencing

DNA was extracted using Magnetic Soil And Stool DNA Kit (Qiagen, Hilden, Germany) following the manufacturer’s protocol. DNA concentration and quality were checked using an Agilent 5400 system and agarose gel electrophoresis. A total of 0.4 μg DNA per sample was used for the DNA library preparations. Then, the library was sequenced on a Novaseq 6000 platform (Illumina, San Diego, CA) adopting a 150-bp paired-end sequencing strategy.

### Curated collection of publicly available samples

We conducted a systematic literature search in May 2024 using Web of Science and Google Scholar with the search terms such as ‘polar,’ ‘permafrost,’ ‘glacier,’ ‘Arctic,’ ‘cold seep,’ ‘hot spring,’ ‘hypersaline,’ ‘saline soil,’ ‘salt lake,’ ‘deep sea,’ ‘deep ocean,’ ‘terrestrial subsurface,’ ‘groundwater,’ ‘hydrothermal plumes,’ ‘hydrothermal vents,’ ‘hydrothermal mats,’ ‘hydrothermal sediment,’ ‘acid mine drainage,’ ‘mining,’ and ‘mine tailing.’ Corresponding metagenomic datasets were retrieved from public repositories such as the NCBI Sequence Read Archive (SRA) and the European Nucleotide Archive (ENA). Environmental metadata were manually curated based on descriptions in papers and databases. Samples with ambiguous origins or those from laboratory enrichment or manipulation were removed. Amplicon sequencing samples and metagenomic samples with low sequencing depth (<200 Mb) were also removed. To ensure platform consistency and minimize technical bias, only datasets generated using the Illumina sequencing platform were retained. We applied additional filtering criteria to refine the dataset: deep sea samples were restricted to those collected at depths >1,000 m, cryosphere samples included only samples from habitats with temperatures <4 °C, saline-alkaline (excluding salt lakes) samples were limited to those with EC >30 dS/m or pH >8.5.

We also collected corresponding environmental parameters across habitats to support downstream analysis. For acid mine samples, pH values were collected; for subsurface, deep sea and hydrothermal plume samples, sampling depths were collected; for cryosphere, hot spring, and hydrothermal plume samples, *in situ* temperature was collected; for saline-alkaline samples, salinity and EC values were collected.

### Metagenomic sequencing data assembly and binning

Both in-house and public metagenomic data were processed using a unified bioinformatic workflow. In detail, raw sequencing reads were quality-filtered with Trim Galore (v0.5.0) (https://github.com/FelixKrueger/TrimGalore) to remove adapters and reads with quality scores <20. *De novo* assembly of clean reads was performed separately for each sample using MEGAHIT (v1.1.3) ^49^ with default settings.

Contigs generated from single-sample assemblies were filtered to retain only those >1,500 bp using Seqtk (v1.5.0) (https://github.com/lh3/seqtk). Clean reads were then mapped back to their corresponding contigs using Bowtie2 (v2.3.5) ^50^. Mapping results were converted into BAM format using SAMtools (v1.9.0), followed by sorting and indexing with the same tool. Metagenomic binning was performed on the filtered contigs using three complementary algorithms: MaxBin2 (v2.2.6) ^51^, MetaBAT2 (v2.12.1) ^52^, and CONCOCT (v1.0.0) ^53^. The resulting bins were integrated using DAS Tool (v1.1.7) ^54^ to produce a non-redundant set of MAGs. The completeness and contamination of all MAGs were assessed using two complementary methods: CheckM1 (v1.2.3; lineage_wf) ^55^ and CheckM2 (v1.0.2; predict workflow) ^56^. MAGs were retained for downstream analyses if both CheckM v1 and CheckM v2 reported a completeness of ≥50% and a contamination of ≤10% according to the MIMAG guidelines ^27^. For high-quality genome identification, Barrnap (v0.9) (https://github.com/tseemann/barrnap) was used to detect ribosomal RNA (rRNA) genes, with the options ‘--kingdom bac’ and ‘--kingdom arc’ applied for bacterial and archaeal MAGs, respectively, along with ‘--reject 0.01’ and ‘--e-value 1e-3’. Transfer RNA (tRNA) genes were annotated using tRNAscan-SE (v2.0.12) ^57^, applying the ‘-A’ mode for archaeal MAGs and ‘-B’ for bacterial MAGs. MAGs were classified as high-quality if they met the MIMAG criteria ^27^: >90% completeness, <5% contamination, ≥18 tRNA genes, and the presence of 5S, 16S, and 23S rRNA genes. MAGs that met only the minimum completeness and contamination thresholds were categorized as medium-quality. The optimal growth temperature was predicted using MetaThermo (v1.0) ^58^ based on genomic sequences, as previously described ^59^.

### Species-level MAG clustering and taxonomic annotation

All MAGs were dereplicated and clustered into 21,805 SGBs at 95% ANI using dRep (v3.4.5) ^60^ with the parameters ‘-pa 0.9 -sa 0.95 -nc 0.3’. For each cluster, a representative genome was chosen based on the quality score. All SGBs were taxonomically classified using GTDB-Tk (v2.4.0) ^61^ with the GTDB (Release R220) (http://gtdb.ecogenomic.org/), and standardized taxonomic labels were assigned.

SGBs with an ANI <0.95 and coverage <0.3 relative to reference genomes were designated as novel (unknown SGBs, uSGBs). SGB consisting of MAGs from the same habitat were defined as habitat-specific, and the corresponding MAGs were habitat-specific. The SGBs consisting of MAGs from different habitats were defined as multi-habitat, and the corresponding MAGs were multi-habitat.

### Estimation of the abundance of kSGBs and uSGBs

To estimate the abundance of known SGBs (kSGBs) and uSGBs, all 1,462 metagenomic samples were mapped against the SGBs using kraken2 (v2.1.3) ^62^. In detail, the taxonomic id of each SGB was defined using a custom python script (gtdb_to_kraken2.py). Then, SGBs were compiled into a single FASTA file and used to construct a custom database with the ‘kraken2-build’ command. The relative abundances of SGBs in each sample were estimated using Kraken2 by mapping clean reads against the custom database.

### Phylogenetic analysis of SGBs

The aligned protein sequences (120 bacterial markers and 53 archaeal markers) were extracted from GTDB-Tk intermediate output files and processed with BMGE (v1.12) ^63^ to retain phylogenetically informative regions. Phylogenetic trees were then constructed using FastTree 2 (v2.1.10) ^64^ with default parameters and visualized and edited using the Interactive Tree Of Life (iTOL) platform (v7.0.0) ^65^.

To quantify the phylogenetic diversity gain introduced by SEGC-derived archaeal uSGBs, we constructed a comprehensive archaeal phylogenetic tree using the same approach as above, comprising 2,654 SGBs from the SEGC dataset and 5,869 archaeal genomes from the GTDB (Release R220). Using GenomeTreeTk (v0.1.8) (https://github.com/dparks1134/GenomeTreeTk), the baseline phylogenetic diversity was first estimated from GTDB archaeal genomes alone. The phylogenetic diversity was then recalculated after incorporating the SEGC-derived uSGBs. The phylogenetic diversity gain was defined as the absolute difference between the two values, and the relative increase was expressed as a percentage of the baseline ^66^. To further uncover the phylogenetic diversity gain introduced by SEGC-derived uSGBs at finer taxonomic resolution, we applied the same approach at the genus level. For each genus containing both previously catalogued and newly recovered species, phylogenetic diversity was calculated before and after incorporating the uSGBs. The absolute and relative phylogenetic diversity gains were then quantified per genus to assess lineage-specific contributions to archaeal diversity.

### Comparison with public microbial genome databases at species-level

To investigate the taxonomic uniqueness of microorganisms inhabiting extreme environments, we compared SGBs from the SEGC against five representative genome catalogues derived from aquatic ^28^, crop ^29^, human ^30^, ocean ^31^, and soil ^32^ microbiomes. The genome collections are described in the Data and materials availability section. MAGs within each catalogue were de-replicated at 95% ANI using dRep (v3.4.5) ^60^ with the parameters ‘-pa 0.9 -sa 0.95 -nc 0.3’, and the highest-quality genome within each cluster was selected as the representative species-level genome for downstream analyses.

For each reference catalogue, sketch indices were generated using skani (v. 0.3.0) ^67^, followed by skani search to identify candidate closest genomes for every SEGC SGB based on approximate ANI. To obtain accurate ANI values, we re-computed ANI for the candidate matches using fastANI (v. 1.33) ^68^ with default parameters except --minFraction 0.3 and --fragLen 1500. For each SGB, only the reference genome with the highest ANI value was retained. SGBs with fastANI >95% to any genome within a given reference catalogue were considered to represent species contained in that catalogue, whereas SGBs that failed to reach this cutoff across all five catalogues were regarded as species absent from conventional microbiomes.

### Analysis of secondary metabolite biosynthetic gene clusters

BGCs for secondary metabolites in the SEGC were identified using antiSMASH (v. 7.0) ^69^. These BGCs were then categorized into eight classes by BiG-SCAPE (v. 2.0) ^70^, including ‘PKSI’, ‘PKS-NPR_Hybrids’, ‘PKSothers’, ‘NRPS’, ‘RiPPs’, ‘Terpene’, ‘Saccharides’ and ‘Others’. The completeness of each BGC was analyzed using BiG-SLiCE (v. 2.0) ^70^ by checking whether it was located on a contig edge, as described in previous studies ^71^. BGCs were clustered into GCFs using BiG-SCAPE based on a default cutoff (c = 0.3). The GCFs consisting of BGCs from the same habitat were defined as habitat-specific, and the corresponding BGCs were habitat-specific. The GCFs consisting of BGCs from different habitats were defined as multi-habitat, and the corresponding BGCs were multi-habitat.

BiG-SLiCE distances were used to assess the novelty of SEGC-derived BGCs by comparing them with BiG-FAM database ^72^, a recently released database containing nearly 1.2 million BGCs. We used a BiG-SLiCE distance >900 as the threshold for novelty, consistent with previous recommendations ^73^. For each GCF, the BiG-SLiCE distance was determined by the minimum distance of its member BGCs.

### Habitat preferences of structural domains

For each phylum within each habitat, the BGCs were de-replicated based on GCF clustering, and the longest BGC was selected as the representative for each GCF. This treatment yielded a non-redundant subset of BGCs for each phylum within every habitat. Protein-coding genes in non-redundant BGCs were predicted with Prodigal (v2.6.3) ^74^ using default settings. The corresponding protein sequences were annotated with structural domains using PfamScan ^75^. For each phylum, the frequencies of each structural domain were quantified within the corresponding BGC subsets. The structural domain preference was assessed by comparing the occurrence ratios of structural domains across habitats using the Kruskal–Wallis test. To avoid sparse artifacts, only BGC subsets containing more than five BGCs were analyzed. False discovery rate (FDR) correction was applied for multiple comparisons. Structural domains with FDR-adjusted *P* values < 0.05 were considered to be non-randomly distributed across habitats, indicating significant habitat-associated preferences.

### Terpene pathway reconstruction and BGC classification

Terpene biosynthetic pathways were reconstructed based on curated information from KEGG (https://www.kegg.jp/) and Metacyc (https://metacyc.org/). Marker genes that are uniquely present in specific terpene biosynthetic pathway were used to distinguish the BGCs of different terpene metabolites. The central pathway of terpene biosynthesis involves the production of the C₅ building blocks isopentenyl diphosphate (IPP) and dimethylallyl diphosphate (DMAPP). Sequential condensation by prenyltransferases yields the precursor diphosphates geranyl diphosphate (GPP, C₁₀), farnesyl diphosphate (FPP, C₁₅), and geranylgeranyl diphosphate (GGPP, C₂₀). Subsequent modification and cyclization of GPP, FPP, and GGPP give rise to distinct monoterpene, diterpene, sesquiterpene, triterpene, and tetraterpene scaffolds, respectively (Fig. 5).

Protein-coding genes in terpene BGCs were predicted with Prodigal (v. 2.6.3) ^74^, and enzyme functions were inferred using CLEAN (v. 1.0.1) ^76^, a contrastive-learning-based enzyme annotation framework, with medium confidence thresholds (confidence ≥0.2) as previously reported ^77^. Predicted EC numbers were used to assign terpene categories (e.g., tetraterpenes), and specific biosynthetic pathways or products (e.g., the retinal biosynthetic pathway) were further identified based on marker genes.

### Geographic distribution and transcriptional activity of the retinal biosynthetic pathway

Retinal biosynthetic genes within terpene BGCs were identified based on EC annotations predicted by CLEAN ^76^. Bacteriorhodopsin-like proteins were detected using Pfam domain annotation with PfamScan ^75^. The geographic distribution was analyzed based on the origins of terpene BGCs with the retinal biosynthetic genes. The transcriptional activity was evaluated using metatranscriptomic sequencing data. In total, 271 cryosphere and 330 saline–alkaline metagenomic samples were analyzed, among which 35 cryosphere samples had corresponding metatranscriptomes. These 35 samples were derived from the Arctic Ocean and were collected as part of the Tara Oceans expedition ^24^.

Raw metatranscriptomic reads were quality-filtered with Trim Galore (v0.5.0) to remove adapters and low-quality reads (Q<20). Retinal pathway genes and bacteriorhodopsin-like protein sequences were indexed using Bowtie2 (v2.4), and clean reads were aligned using default sensitive parameters. Alignment files were converted, sorted, and indexed with SAMtools. Gene-level expression was quantified as FPKM (Fragments Per Kilobase per Million mapped reads), normalized by total raw read counts per sample.

### Statistical analysis

All comparisons between two groups were assessed using the Wilcoxon rank-sum test. Effect sizes were calculated using Cohen’s *d*, and absolute values (|*d*|) were reported to quantify the magnitude of group differences. All statistical analyses were performed using R (v. 4.0.0).

## Data and materials availability

The raw data of metagenomic sequencing of 115 soil samples were deposited in the NCBI database under the accession numbers PRJNA1285087. The accessions of the metagenomic and metatranscriptomic data from public repositories were provided in Supplementary materials. The MAG’s availability of aquatic ^28^, crop ^29^, human ^30^, ocean ^31^, and soil ^32^ habitats were listed in Supplementary materials. The fasta files of all MAGs and gbk files of BGCs were uploaded into the Zenodo (https://doi.org/10.5281/zenodo.15788452). The code developed in this study has been made available on GitHub (https://github.com/durubing-jn/SEGC).

## Acknowledgments

Funding

This work was supported by Basic Research Program of Jiangsu (BK20250037), the Project of National Natural Science Foundation of China (22378210), the Natural Science Foundation of the Jiangsu Higher Education Institutions of China (23KJA180003), Project of Fund for Stable Support to Agricul-tural Sci-Tech Renovation (xjnkywdzc-2024001), Postdoctoral Fellow-ship Program of CPSF (GZC20240728), the Jiangsu Funding Program for Excellent Postdoctoral Talent (2024ZB271), the Jiangsu Basic Research Center for Synthetic Biology Grant (No. BK20233003), State Key Laboratory of Microbial Technology Open Project Fund (Project NO. M2025-15).

